## Supplementary material for "Mitochondrial complex I deficiency occurs in skeletal muscle of a subgroup of individuals with Parkinson’s disease": Inventory of Supporting Information

**Supplementary Figure 1.**
Validation of quadruple immunohistochemistry for quantitative MRC complex assessment.

**Supplementary Figure 2.**
Examples of cytochrome c oxidase/succinate dehydrogenase histochemistry.

**Supplementary Figure 3.**
Measurement of immunohistochemistry fluorescence signal in individual muscle fibers and large section areas.

**Supplementary Figure 4.**
Correlation between immunohistochemistry fluorescence measurement in multiple single fibers and in a large section area.

**Supplementary Figure 5.**
Immunohistochemistry of complexes I and IV in PD single muscle fibers without adjusting for staining batch.

**Supplementary Figure 6.**
Mitochondrial content in single muscle fibers.

**Supplementary Figure 7.**
Spectrophotometric activity measurement in muscle without adjusting for batch effects.

**Supplementary Figure 8.**
Spectrophotometric activity measurement in muscle including smokers and adjusting for batch effects.

**Supplementary Figure 9.**
Spectrophotometric activity measurement in muscle including smokers and without adjusting for batch effects.

**Supplementary Figure 10.**
Power as a function of sample size for complex I activity data.

**Supplementary Figure 11.**
Selection of samples for mtDNA analysis in single muscle fibers.

**Supplementary Figure 12.**
Selection of samples for mtDNA analysis in bulk muscle tissue.

**Supplementary Figure 13.**
Separation of individual muscle fibers based on laminin staining.

**Supplementary Table 1.**
Demographic and clinical characteristics per analysis.

**Supplementary Table 2.**
Comparison of COX/SDH histochemistry results in the PD and control groups.

**Supplementary Table 3.**
Linear mixed effects model of VDAC1 fluorescence intensity level in single muscle fibers.

**Supplementary Table 4.**
Linear regression models of complex I and IV level and VDAC1 fluorescence level in large muscle sections areas.

**Supplementary Table 5a.**
Linear regression models of enzymatic activity of CI, II, III, IV and citrate synthase in individuals with PD and controls with coefficients for batch variables included in the table.

**Supplementary Table 5b.**
Linear regression models of enzymatic activity of CI, II, III, IV and citrate synthase in individuals with PD and controls with smokers included.

**Supplementary Table 6.**
Linear regression model of enzymatic activity of CI in individuals with PD.

**Supplementary Table 7.**
Comparison of clinical and demographic characteristics between the PD subgroups.

**Supplementary Table 8.**
Linear mixed effects model of complex I level in single muscle fibers in individuals with PD and controls.

**Supplementary Table 9**.
ANOVA comparison evaluating the impact of complex I quantity on complex I activity.

**Supplementary Table 10.**
Linear regression model of enzymatic activity of CI in individuals with PD and controls.

**Supplementary Table 11.**
Linear mixed effects model of mtDNA copy number in single muscle fibers.

**Supplementary Table 12.**
Linear mixed effects model of mtDNA major arc deletion levels in single muscle fibers.

**Supplementary Table 13.**
Linear mixed effects model of heteroplasmic load in single muscle fibers.

**Supplementary Table 14.**
Wilcoxon rank sum tests of mtDNA data in bulk muscle tissue.

**Supplementary Table 15.**
Linear regression model of heteroplasmic load in bulk muscle tissue as a function of CI activity and enzymatic activity measurement batch.

**Supplementary Table 16.**
Inclusion and exclusion criteria for the STRAT-PARK cohort.

**Supplementary Table 17.**
Inclusion and exclusion criteria for the NADPARK study.

**Supplementary Table 18.**
Inclusion and exclusion criteria for the STRAT-COG study.

**Supplementary Table 19.**
Olympus VS120 fluorescence filters.

**Supplementary Data 1.**
Experimental allocation of subjects.

**Supplementary Data 2.**
COX/SDH histochemistry data.

**Supplementary Data 3.**
Single muscle fiber IHC data.

**Supplementary Data 4.**
Large muscle section area IHC data.

**Supplementary Data 5.**
Muscle MRC activity data.

**Supplementary Data 6.**
Single muscle fiber copy number and deletion level data.

**Supplementary Data 7.**
Single muscle fiber heteroplasmic load data.

**Supplementary Data 8.**
Single muscle fiber heteroplasmic load distribution data.

**Supplementary Data 9.**
Bulk muscle tissue copy number and deletion level data.

**Supplementary Data 10.**
Bulk muscle tissue heteroplasmic load data.

**Supplementary Data 11.**
Bulk muscle tissue heteroplasmic load distribution data.

**Supplementary Data 12.**
Muscle IHC and COX/SDH data from an individual with a single mitochondrial DNA deletion.
