## Supplementary information for "Mitochondrial complex I deficiency occurs in skeletal muscle of a subgroup of individuals with Parkinson’s disease"

Simon Ulvenes Kverneng *et al.*

Corresponding author: Charalampos Tzoulis

**This PDF file includes:**

Supplementary Figs. 1 to 13

Supplementary Tables 1 to 19

**Other Supplementary Information for this manuscript include the following (separate files):**

Supplementary Data 1 to 12

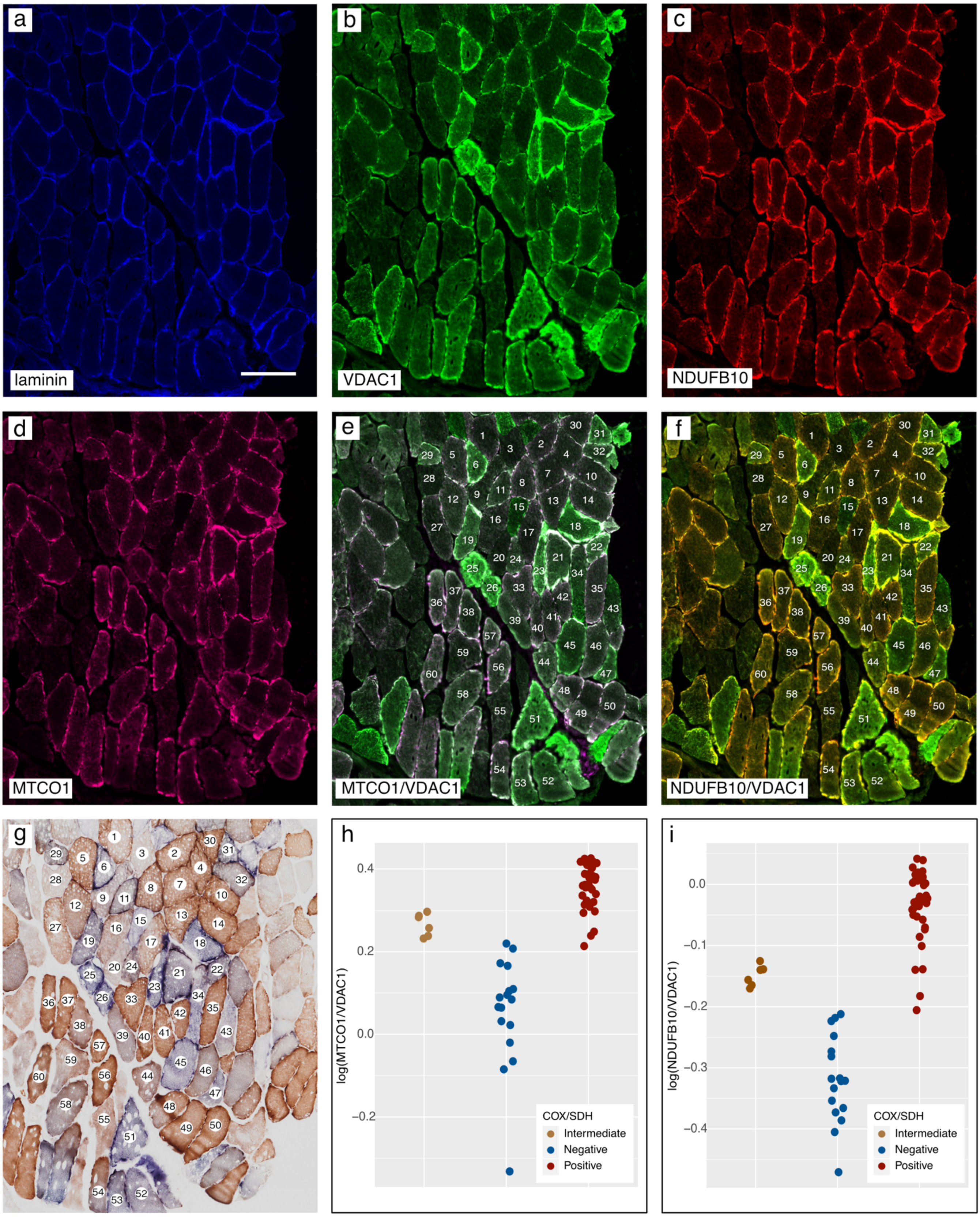
a
laminin
b
VDAC1
c
NDUFB10
d
MTCO1
e
MTCO1/VDAC1
f
NDUFB10/VDAC1
g
h
log(MTCO1/VDAC1)
COX/SDH
Intermediate
Negative
Positive
i
log(NDUFB10/VDAC1)
COX/SDH
Intermediate
Negative
Positive

**Supplementary Figure 1. Validation of quadruple immunohistochemistry for quantitative MRC complex assessment.**

Quadruple immunohistochemistry and cytochrome c oxidase/succinate dehydrogenase (COX/SDH) histochemical staining of two consecutive sections from a skeletal muscle biopsy harboring a single mtDNA deletion. Immunohistochemistry staining of laminin **(a)**, scale bar: 200 µm, the mitochondrial marker porin (VDAC1;**b**), subunit NDUFB10 of complex I (CI; **c**), subunit MTCO1 of complex IV (CIV, **d**). **(e)** Merged MTCO1 and VDAC1 signal. **(f)** Merged NDUFB10 and VDAC1 signal. **(g)** Consecutive section from the same skeletal muscle as in a-f stained with COX/SDH histochemistry. COX-status was qualitatively assessed (“positive”, “intermediate” or “negative”) by two readers (SUK and CT). 60 corresponding muscle fibers are labeled in e, f and g. **(h)** VDAC1-normalized CIV fluorescence intensity (log(MTCO1/VDAC1)) compared to COX/SDH status in the 60 labeled muscle fibers. **(i)** VDAC1-normalised CI fluorescence intensity (log(NDUFB10/VDAC1)) compared to COX/SDH status in the 60 labeled muscle fibers. All fluorescence intensities were adjusted for background fluorescence.

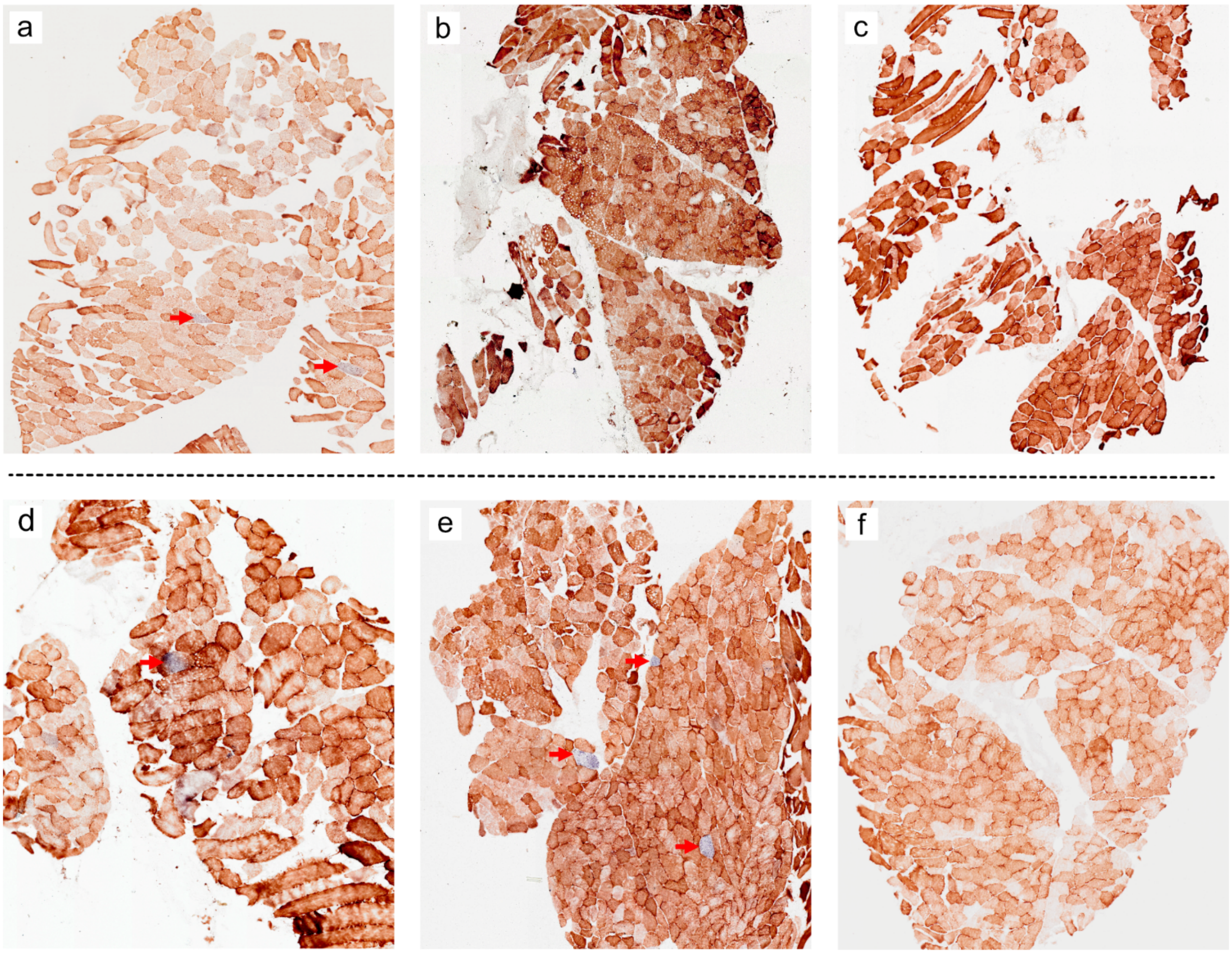


**Supplementary Figure 2. Cytochrome c oxidase/succinate dehydrogenase histochemistry.** Representative examples of cytochrome c oxidase/succinate dehydrogenase (COX/SDH) histochemical staining from muscle biopsies of three neurologically healthy controls (a – c) and three individuals with Parkinson's disease (d – f). COX-positive fibers stain brown, while COX-negative and SDH-positive fibers stain blue (red arrows).

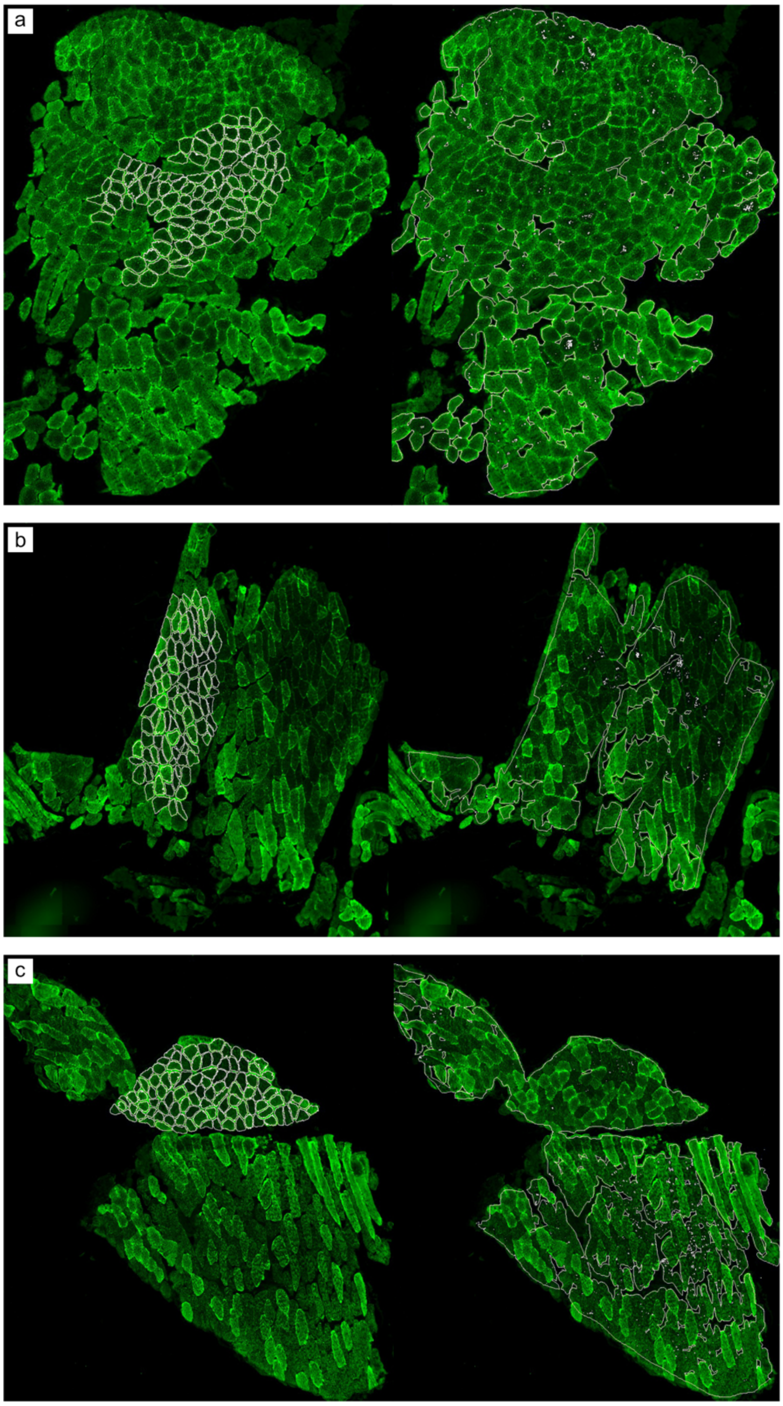


**Supplementary Figure 3. Measurement of immunohistochemistry fluorescence signal in individual muscle fibers and large section areas.**

**(a-c)** Representative examples of regions of interest (ROIs) used for measuring immunohistochemistry fluorescence signal in individual muscle fibers (left image) and large section areas encompassing the majority of muscle fibers (right image) from the same sections.

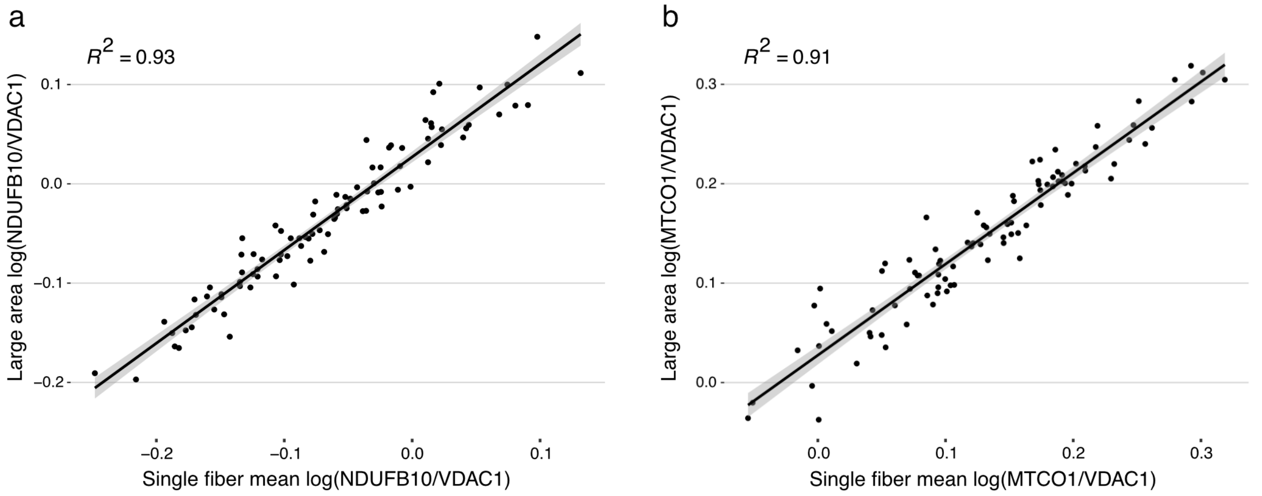


**Supplementary Figure 4. Correlation between immunohistochemistry fluorescence measurement in multiple single fibers and in a large section area.**

In each section, fluorescence intensity was measured in multiple single muscle fibers ($n$ = 75 - 100 per section) and in a single large area covering the majority of fibers of the section. **(a)** Correlation between the mean of single fiber CI levels and CI level in a single large area from the same section. Individuals with PD and control individuals are plotted together. Each dot represents one individual. **(b)** Correlation between the mean of single fiber CIV levels and CIV level in a single large area from the same section. Individuals with PD and control individuals are plotted together. Each dot represents one individual.

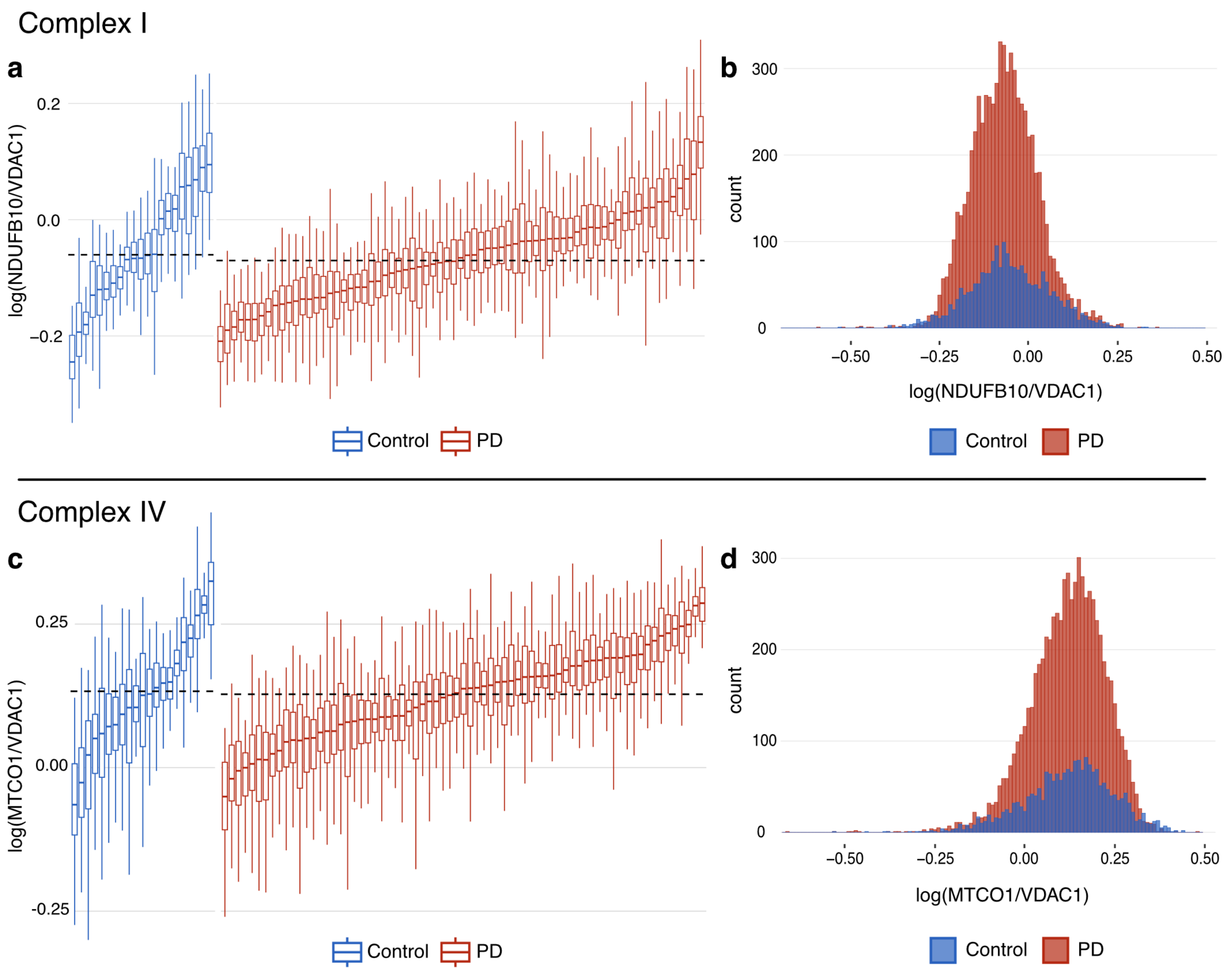


**Supplementary Figure 5. Immunohistochemistry of complexes I and IV in PD single muscle fibers without adjusting for staining batch.**

Complex I (NDUFB10) and complex IV (MTCO1) fluorescence intensity normalized to mitochondrial mass (VDAC1) in single muscle fibers (*n* = 75-100 per individual) in the PD (red) and control (blue) groups. Values are log transformed. Data have not been adjusted for staining batch. Boxplots **(a, c)** show individual-level distributions of single fiber measurement where each box represents one individual. Boxes: median and interquartile range (IQR); whiskers: 1.5 x IQR from the lower and upper quartiles. Individuals are sorted by median values from left to right. Dashed lines show the group-level medians of the PD and control groups. The histograms **(b, d)** represent group-level distributions of single fiber measurement in the PD and control groups.

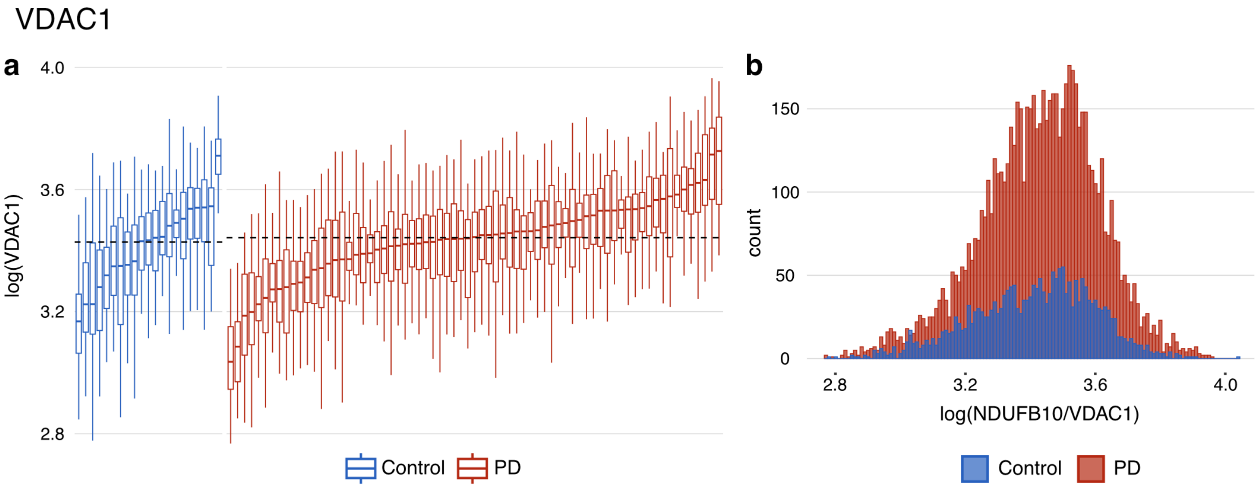


**Supplementary Figure 6. Mitochondrial content in single muscle fibers.**

VDAC1-detection in single fibers ($n$ = 75-100 per individual) in the PD (red) and control (blue) groups. Values are log transformed. For the purpose of visualization, the data have been adjusted for the effect of immunohistochemistry staining batch by regressing out this variable (see Methods section). The boxplot **(a)** shows individual-level distributions of single fiber measurement where each box represents one individual. Boxes: median and interquartile range (IQR); whiskers: 1.5 x IQR from the lower and upper quartiles. Individuals are sorted by median values from left to right. Dashed lines show the group-level medians of the PD and control groups. The histogram **(b)** represents group-level distribution of single fiber measurements in the PD and control groups.

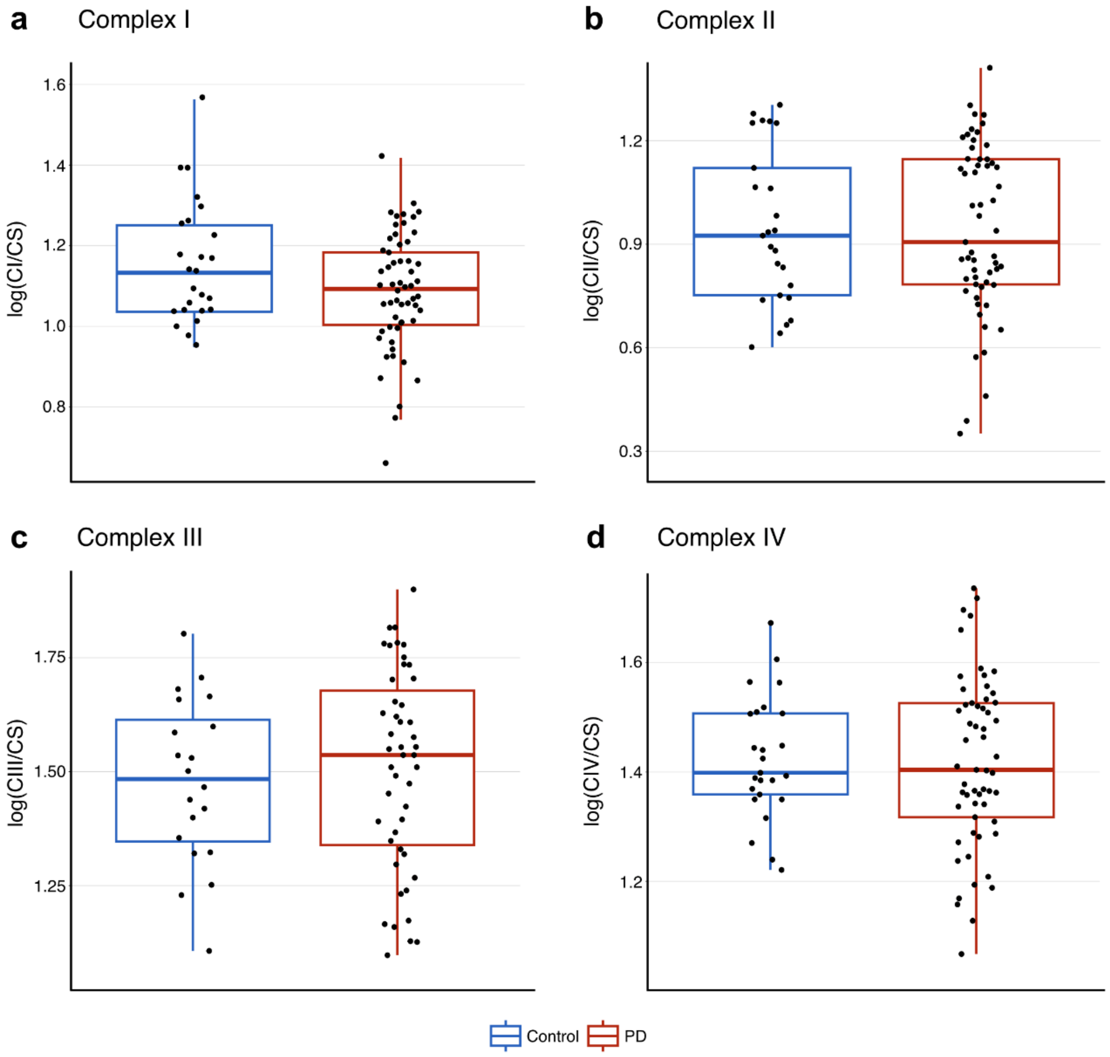


**Supplementary Figure 7. Spectrophotometric activity measurement in muscle without adjusting for batch effects.**

Activities of complexes I-IV (CI-CIV), normalized to citrate synthase (CS) activity. Values are log transformed. Smokers have been removed. Red boxplots represent the PD group, and blue boxplots represent the control group. Boxes: median and interquartile range (IQR); whiskers: 1.5 x IQR from the lower and upper quartiles. Each dot represents one individual. **(a)** CS-normalized CI activity. **(b)** CS-normalized CII activity. **(c)** CS-normalized CIII activity. Measurements from 15 samples were excluded due to technical issues with the reduction of decylubiquinone (Supplementary data 1). **(d)** CS-normalized CIV activity.

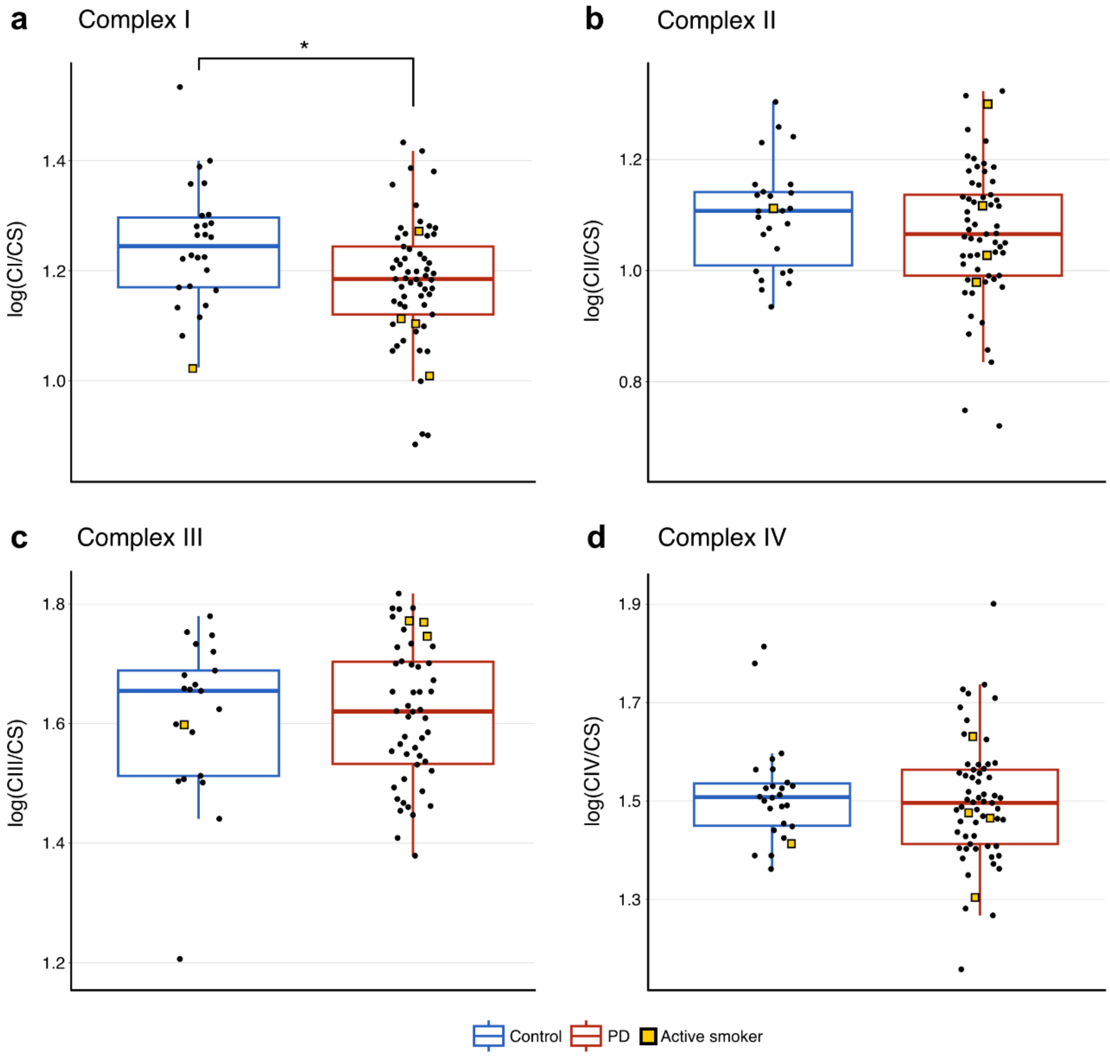


**Supplementary Figure 8. Spectrophotometric activity measurement in muscle including smokers and adjusting for batch effects.**

Activities of complexes I-IV (CI-CIV), normalized to citrate synthase (CS) activity. Smokers are included (yellow squares). Values are log transformed. The data have been adjusted for the effect of measurement batch by regressing out this variable (see Methods section). Red boxplots represent the PD group, and blue boxplots represent the control group. Boxes: median and interquartile range (IQR); whiskers: 1.5 x IQR from the lower and upper quartiles. Each dot represents one individual. **(a)** CS-normalized CI activity. **(b)** CS-normalized CII activity. **(c)** CS-normalized CIII activity. 16 samples were excluded due to technical issues with the reduction of decylubiquinone (Supplementary Data 1). **(d)** CS-normalized CIV activity.

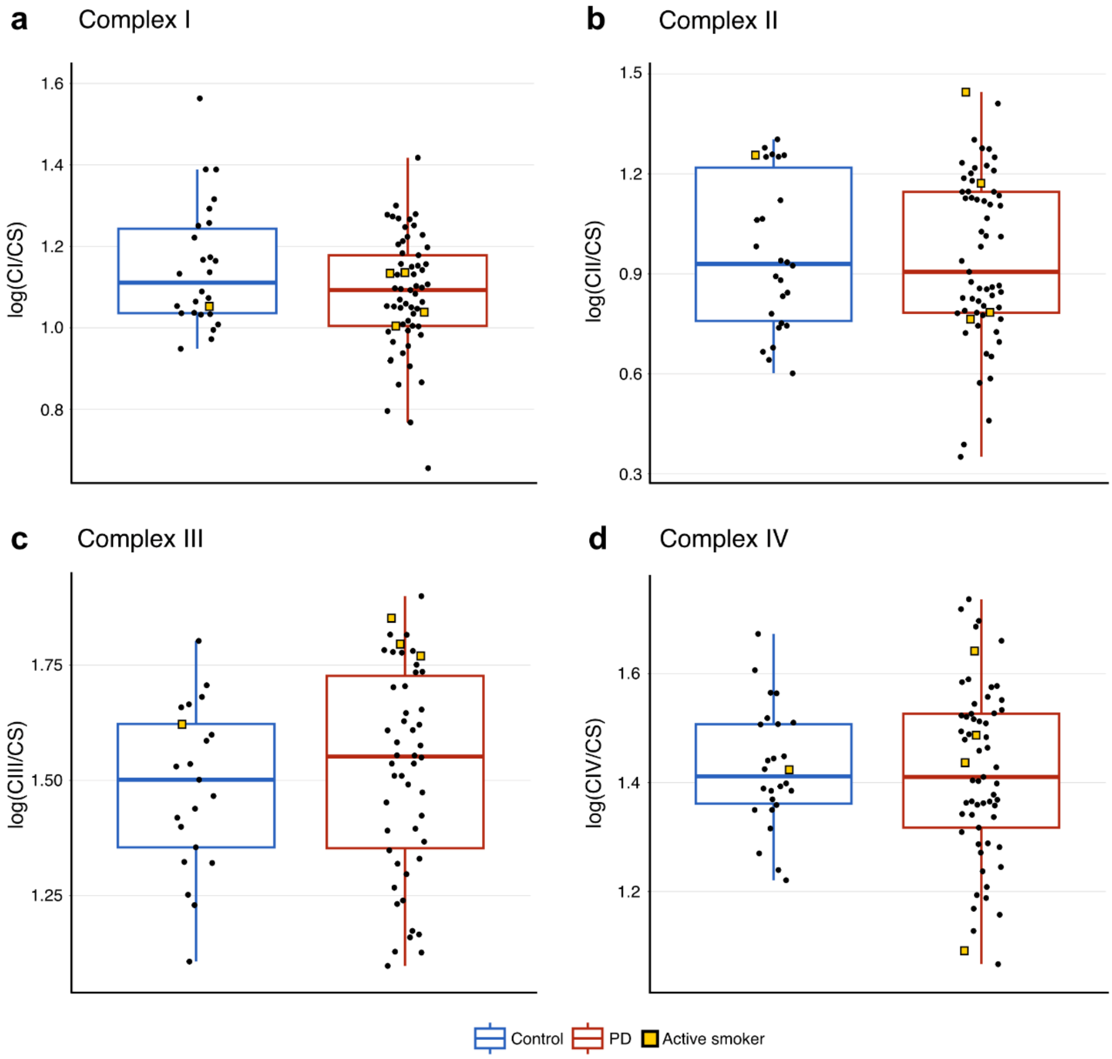


**Supplementary Figure 9. Spectrophotometric activity measurement in muscle including smokers and without adjusting for batch effects.**

Activities of complexes I-IV (CI-CIV), normalized to citrate synthase (CS) activity. Values are log transformed. Smokers (yellow squares) are included. Red boxplots represent the PD group, and blue boxplots represent the control group. Boxes: median and interquartile range (IQR); whiskers: 1.5 x IQR from the lower and upper quartiles. Each dot represents one individual. **(a)** CS-normalized CI activity. **(b)** CS-normalized CII activity. **(c)** CS-normalized CIII activity. 16 samples were excluded from this analysis due to technical issues with the reduction of decylubiquinone (Supplementary Data 1). **(d)** CS-normalized CIV activity.

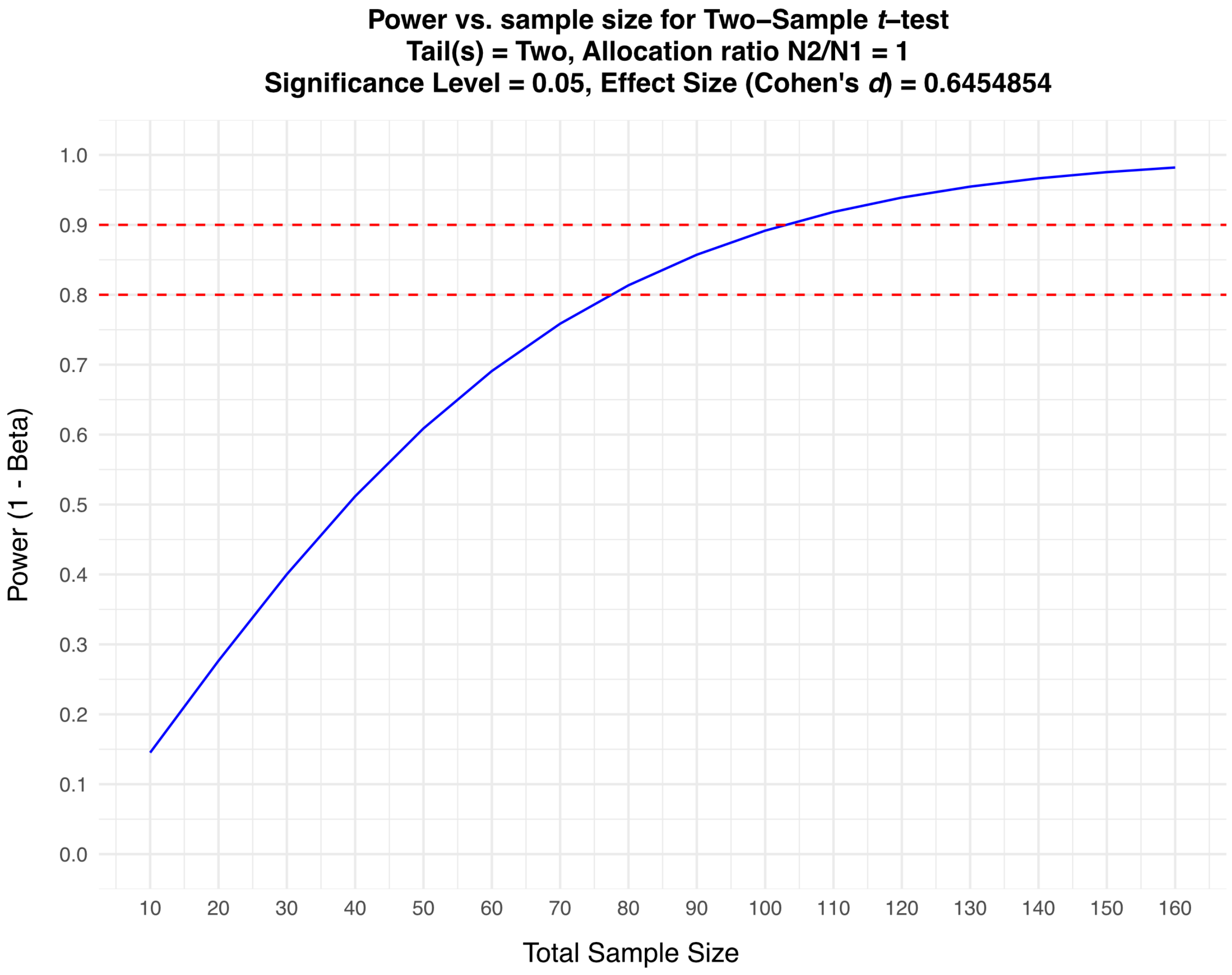


**Supplementary Figure 10. Power as a function of sample size for complex I activity data.** After adjusting the data for the effect of measurement batch by regressing out this variable (see Methods section), the difference in citrate synthase-normalized complex I activity (CI/CS) between the patient and control groups corresponded to a medium effect size (Cohen’s *d* of 0.65). The plot shows the total sample size needed for different power values for a two-sample *t*-test given this effect size. Dashed red lines indicate 80% and 90% power.

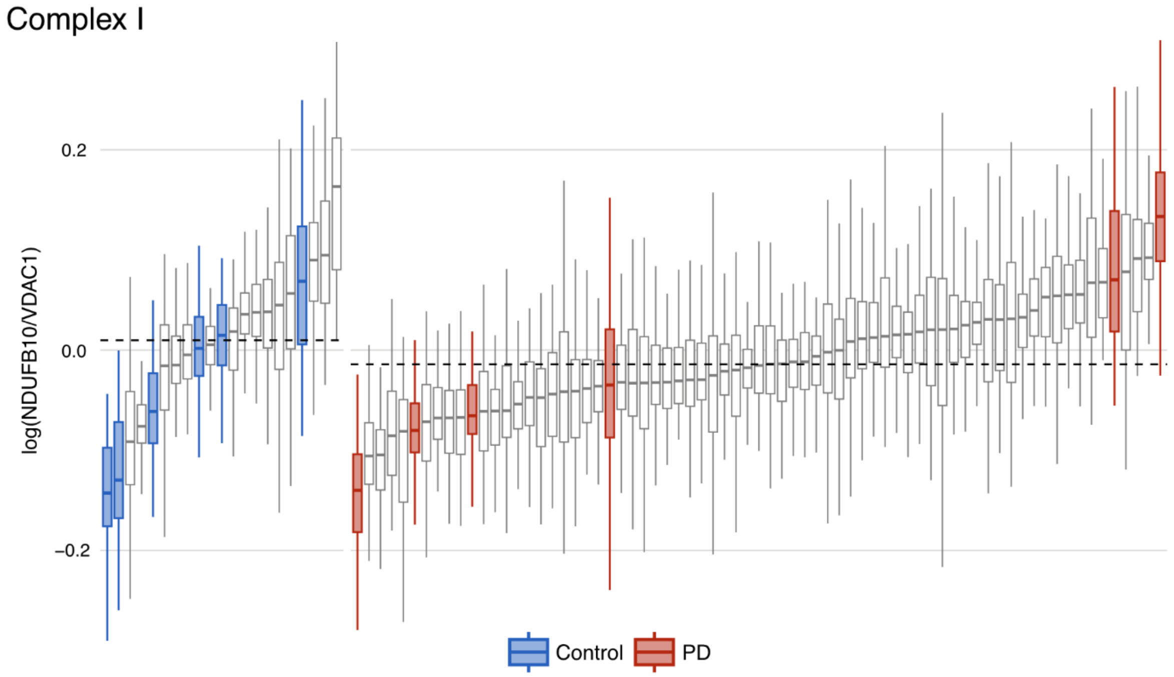


**Supplementary Figure 11. Selection of samples for mtDNA analysis in single muscle fibers.** Six PD samples (red) and six control samples (blue), spanning the range of median complex I quantity, were selected for laser-microdissection of multiple single muscle fibers for mtDNA analysis. Boxplots show complex I (NDUFB10) fluorescence intensity normalized to mitochondrial mass (VDAC1) in single muscle fibers (*n* = 75-100 per individual) in the patient and control group. Each boxplot represents one individual. Boxes: median and interquartile range (IQR); whiskers: 1.5 x IQR from the lower and upper quartiles. Individuals are sorted by median values from left to right. Dashed lines represent group-level medians of the entire PD (*n* = 71) and control (*n* = 21) group. Values are log transformed. For the purpose of visualization, the data have been adjusted for the effect of immunohistochemistry staining batch by regressing out this variable (see Methods section).

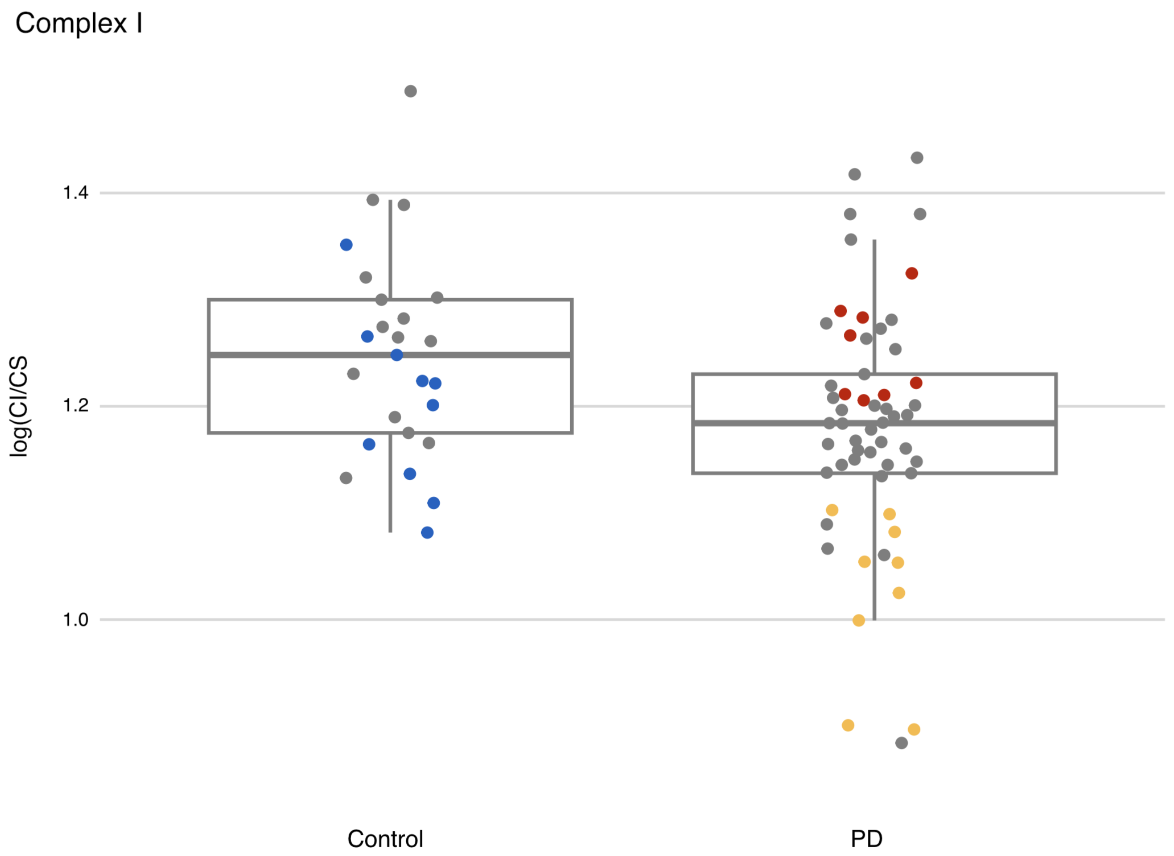


**Supplementary Figure 12. Selection of samples for mtDNA analysis in bulk muscle tissue.** Spectrophotometric activity measurements of complex I (CI) normalized to citrate synthase (CS) in muscle biopsies from individuals with PD and controls are shown. Smokers are not included. 8 PD samples with CS normalized CI activity similar to controls (red), 9 PD samples with low activity levels (yellow), and 10 control samples (blue) were selected for mtDNA analysis in bulk muscle tissue. The three groups were matched for age (Supplementary Table 1). Values are log transformed. The data have been adjusted for the effect of measurement batch by regressing out this variable (see Methods section). Boxes: median and interquartile range (IQR); whiskers: 1.5 x IQR from the lower and upper quartiles. Each dot represents one individual.

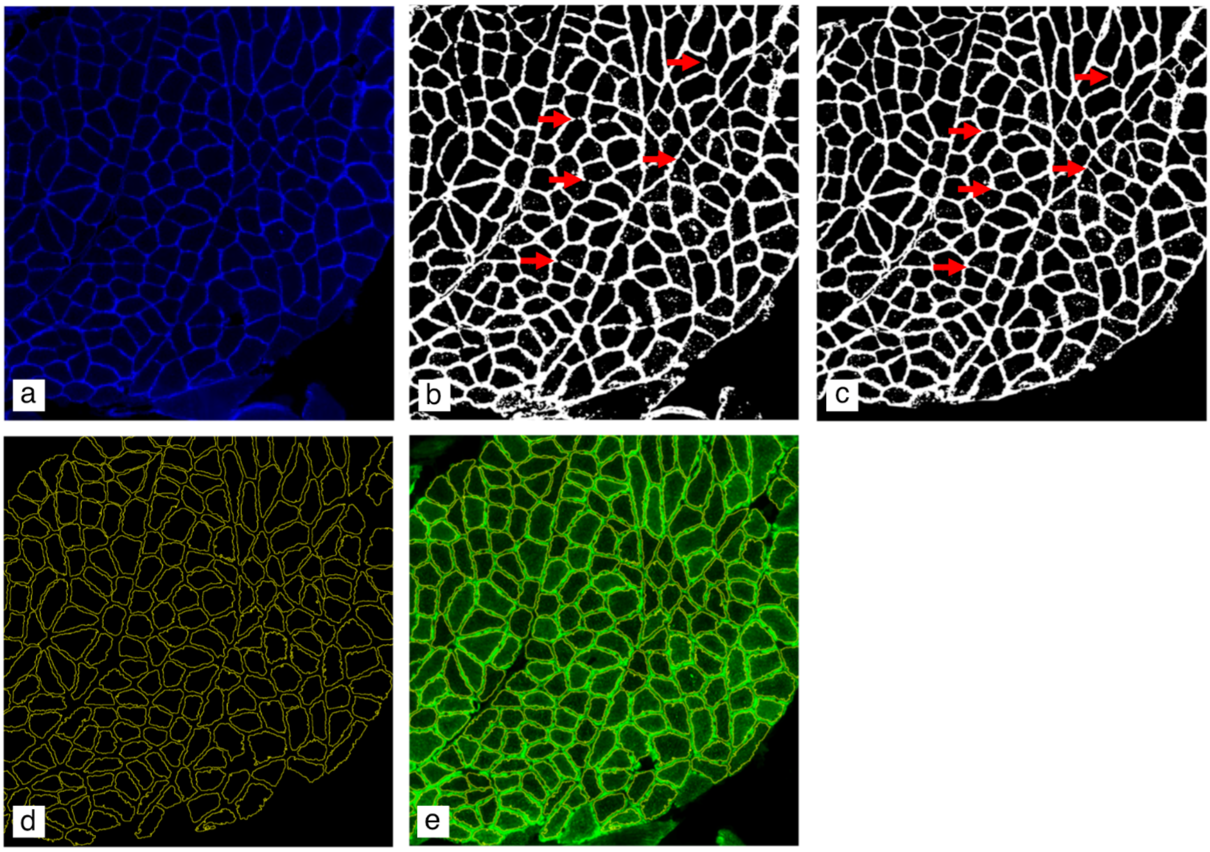


**Supplementary Figure 13. Separation of individual muscle fibers based on laminin staining.** **(a)** Immunohistochemistry detecting laminin. **(b)** The image is processed in ImageJ2 (version 2.3.0/1.53f) to enhance edges and blurring is added to make lines meet. The image is then converted to 8-bit and a threshold is applied and manually adjusted to make a suitable mask. **(c)** Holes in the mask are manually closed using the pencil tool (examples indicated by red arrows). **(d)** Individual regions of interest (ROIs) for each muscle fiber are created. **(e)** ROIs are used to measure fluorescence signal of VDAC1, NDUFB10 and MTCO1 for individual muscle fibers (VDAC1-signal is shown in the image as example).

Supplementary Table 1: Demographic and clinical characteristics per analysis

| Method | *n* | Age | Sex (M : F) | Disease duration in years |
|---|---|---|---|---|
| **COX/SDH histochemistry** | | | | |
| PD | 68 | 65.9 (± 7.8) | 46 : 22 | 5.2 (± 4.4) |
| Control | 21 | 61.1 (± 9.9) | 5 : 16 | |
| **Quadruple immunohistochemistry** | | | | |
| PD | 71 | 66.2 (± 7.8) | 47 : 24 | 5.2 (± 4.3) |
| Control | 21 | 61.1 (± 9.9) | 5 : 16 | |
| **Enzymatic activity (without smokers)** | | | | |
| PD | 57 | 67.1 (± 7.4) | 35 : 22 | 6.5 (± 4.9) |
| Control | 25 | 65.4 (± 12.0) | 8 : 17 | |
| **Enzymatic activity (with smokers)** | | | | |
| PD | 61 | 66.8 (± 7.4) | 37 : 24 | 6.5 (± 4.8) |
| Control | 26 | 65.2 (± 11.8) | 8 : 18 | |
| **mtDNA analyses (bulk tissue)** | | | | |
| PD (normal CI activity) | 8 | 65.0 (± 6.7) | 6 : 2 | 5.2 (± 1.7) |
| PD (low CI activity) | 9 | 66.9 (± 10.9) | 3 : 6 | 5.4 (± 2.7) |
| Control | 10 | 64.9 (± 8.9) | 3 : 7 | |
| **mtDNA analyses (single fibers)** | | | | |
| PD | 6 | 66.8 (± 6.5) | 5 : 1 | 5.1 (± 4.1) |
| Control | 6 | 64.8 (± 9.5) | 1 : 5 | |

Age and disease duration are presented as mean ± standard deviation.

Supplementary Table 2. Comparison of COX/SDH histochemistry results in the PD and control groups

| Variable | Group | Median (IQR) | Test | Test Statistic | *df* | *P*-value |
|---|---|---|---|---|---|---|
| **Proportion of COX negative or intermediate muscle fibers per section** | PD (*n* = 68) | 0 (0.004) | Wilcoxon rank-sum test | *W* = 764 | NA | 0.584 |
| | Control (*n* = 21) | 0 (0.002) | | | | |
| **Number of individuals with any COX negative or intermediate fibers** | | | Pearson's Chi-squared Test (with Yates' continuity correction) | $\chi^2 = 0.07$ | 1 | 0.788 |
| | Any negative or intermediate | No negative or intermediate | | | | |
| PD (*n* = 68) | 27 | 41 | | | | |
| Control (*n* = 21) | 7 | 14 | | | | |

Abbreviations: COX, cytochrome c oxidase; SDH, succinate dehydrogenase; IQR, interquartile range; *df,* degrees of freedom. Significant *P*-values are in bold. Nominal *P*-values are given.

Supplementary Table 3: Linear mixed effects model of VDAC1 fluorescence intensity level in single muscle fibers

| | | *Dependent variable* | |
|---|---|---|---|
| | | **log(VDAC1)** | |
| Predictors | *B* | 95% CI | *P*-value |
| Disease state (PD) | 0.003 | -0.068 – 0.074 | 0.933 |
| Age | 1.3e-04 | -0.003 – 0.004 | 0.942 |
| Sex (Male) | 0.028 | -0.032 – 0.089 | 0.362 |
| Smoking | -0.099 | -0.221 – 0.024 | 0.115 |
| Batch (Batch 2)[a] | 0.188 | 0.132 – 0.244 | **<0.001** |
| **Random Effects** | | | |
| $\sigma^2$ | 0.019 | | |
| $\tau_{00\ \mathrm{Subject}}$ | 0.018 | | |
| ICC | 0.487 | | |
| $n_{\mathrm{Subject}}$ | 92 | | |
| Observations | 9073 | | |
| Marginal $R^2$ / Conditional $R^2$ | 0.191 / 0.585 | | |

Abbreviations: log(VDAC1), measure of mitochondrial content; *B*, regression coefficient (unstandardized); 95% CI, 95% confidence interval of the regression coefficient; $\sigma^2$, residual variance; $\tau_{00\ \mathrm{Subject}}$, random intercept variance; ICC, intraclass correlation coefficient, representing the proportion of total variance in the dependent variable attributable to the grouping structure (i.e, subjects); $n_{\mathrm{Subject}}$, number of study subjects.
Significant *P*-values are in bold. Nominal *P*-values are given.

[a]Immunohistochemistry staining was performed in two batches.

Supplementary Table 4: Linear regression models of complex I and IV level and VDAC1 fluorescence level in large muscle sections areas

| | *Dependent variable* | | | | | | | | |
|---|---|---|---|---|---|---|---|---|---|
| | **log(NDUFB10/VDAC1)** | | | **log(MTCO1/VDAC1)** | | | **log(VDAC1)** | | |
| Predictors | *B* | 95% CI | *P*-value | *B* | 95% CI | *P*-value | *B* | 95% CI | *P*-value |
| Disease state (PD) | -0.016 | -0.046 – 0.014 | 0.285 | -0.016 | -0.056 – 0.023 | 0.412 | -0.016 | -0.089 – 0.057 | 0.668 |
| Age | -0.002 | -0.003 – -2.5e-04 | **0.022** | -0.002 | -0.004 – -3.6e-04 | **0.020** | 9.0e-05 | -0.003 – 0.004 | 0.959 |
| Sex (Male) | 0.017 | -0.008 – 0.043 | 0.182 | 0.047 | 0.013 – 0.081 | **0.007** | 0.010 | -0.052 – 0.073 | 0.746 |
| Smoking | -0.002 | -0.054 – 0.050 | 0.939 | -0.007 | -0.076 – 0.061 | 0.829 | -0.098 | -0.224 – 0.028 | 0.125 |
| Batch (Batch 2)[a] | -0.099 | -0.123 – -0.076 | **<0.001** | 0.052 | 0.021 – 0.083 | **0.001** | 0.216 | 0.159 – 0.274 | **<0.001** |
| Observations | 92 | | | 92 | | | 92 | | |
| $R^2$ / $R^2$ adjusted | 0.495 / 0.465 | | | 0.190 / 0.143 | | | 0.399 / 0.364 | | |

Abbreviations: log(NDUFB10/VDAC1), complex I level adjusted for mitochondrial content; log(MTCO1/VDAC1), complex IV level adjusted for mitochondrial content; log(VDAC1), measure of mitochondrial content; *B*, regression coefficient (unstandardized); 95% CI, 95% confidence interval of the regression coefficient.
Significant *P*-values are in bold. Nominal *P*-values are given.

[a]Immunohistochemistry staining was performed in two batches.

Supplementary Table 5a: Linear regression models of enzymatic activity of CI, II, III, IV and citrate synthase in individuals with PD and controls

| | **log(CI/CS)** | | | **log(CII/CS)** | | | **log(CIII/CS)** | | | **log(CIV/CS)** | | | **log(CS)** | | |
|---|---|---|---|---|---|---|---|---|---|---|---|---|---|---|---|
| Predictors | *B* | 95% CI | *P*-value (adjusted)[a] | *B* | 95% CI | *P*-value (adjusted)[a] | *B* | 95% CI | *P*-value (adjusted)[a] | *B* | 95% CI | *P*-value (adjusted)[a] | *B* | 95% CI | *P*-value |
| Disease state (PD) | -0.079 | -0.137 – -0.021 | **0.008 (0.032)** | -0.043 | -0.103 – 0.018 | 0.163 (0.313) | -0.038 | -0.102 – 0.025 | 0.235 (0.313) | -0.027 | -0.089 – 0.035 | 0.383 (0.383) | 0.034 | -0.090 – 0.159 | 0.582 |
| Age | -0.001 | -0.004 – 0.003 | 0.693 | 0.002 | -0.001 – 0.006 | 0.143 | 0.008 | 0.005 – 0.011 | **<0.001** | 0.003 | -1.8e-04 – 0.007 | 0.063 | -0.001 | -0.008 – 0.006 | 0.826 |
| Sex (Male) | 0.027 | -0.027 – 0.082 | 0.323 | 0.013 | -0.043 – 0.070 | 0.640 | 0.006 | -0.052 – 0.064 | 0.846 | 0.017 | -0.041 – 0.076 | 0.555 | -0.023 | -0.140 – 0.093 | 0.690 |
| Batch 1 | Reference | - | - | - | - | - | - | - | - | - | - | - | - | - | - |
| Batch 2 | 0.067 | -0.029 – 0.163 | 0.171 | 0.158 | 0.058 – 0.258 | **0.002** | 0.095 | 0.002 – 0.188 | **0.045** | 0.049 | -0.053 – 0.152 | 0.341 | -0.027 | -0.233 – 0.180 | 0.797 |
| Batch 3 | -0.125 | -0.216 – -0.035 | **0.007** | 0.115 | 0.021 – 0.209 | **0.017** | -0.239 | -0.326 – -0.151 | **<0.001** | -0.100 | -0.197 – -0.004 | **0.042** | -0.111 | -0.305 – 0.084 | 0.260 |
| Batch 4 | -0.237 | -0.321 – -0.153 | **<0.001** | -0.392 | -0.479 – -0.305 | **<0.001** | -0.307 | -0.387 – -0.228 | **<0.001** | -0.131 | -0.221 – -0.042 | **0.005** | 0.373 | 0.193 – 0.553 | **<0.001** |
| Batch 5 | -0.104 | -0.191 – -0.018 | **0.018** | -0.204 | -0.293 – -0.115 | **<0.001** | | | | -0.195 | -0.287 – -0.103 | **<0.001** | 0.182 | -0.003 – 0.367 | 0.053 |
| Batch 6 | -0.127 | -0.215 – -0.039 | **0.005** | -0.324 | -0.416 – -0.233 | **<0.001** | 0.089 | 0.005 – 0.172 | **0.038** | -0.018 | -0.112 – 0.076 | 0.703 | 0.267 | 0.077 – 0.456 | **0.006** |
| Observations | 82 | | | 82 | | | 67[b] | | | 82 | | | 82 | | |
| $R^2$ / $R^2$ adjusted | 0.481 / 0.424 | | | 0.777 / 0.752 | | | 0.762 / 0.734 | | | 0.348 / 0.276 | | | 0.363 / 0.293 | | |

Abbreviations: CI, complex I activity; CII, complex II activity; CIII, complex III activity; CIV, complex IV activity; CS, citrate synthase activity; x/CS, activity x (CI, CII, CIII, or CIV) normalized to citrate synthase activity; *B*, regression coefficient (unstandardized); 95% CI, 95% confidence interval of the regression coefficient. Significant *P*-values are in bold. Nominal and adjusted *P*-values are given.

[a]Parentheses show *P*-values adjusted for multiple testing using the Benjamini-Hochberg procedure for four tests, i.e., CI/CS, CII/CS, CIII/CS and CIV/CS between the PD and control groups.
[b]Complex III activity measurements from 15 individuals were excluded from the analysis due to technical issues with the reduction of decylubiquinone.

Supplementary Table 5b: Linear regression models of enzymatic activity of CI, II, III, IV and citrate synthase in individuals with PD and controls with smokers included

| | **log(CI/CS)** | | | **log(CII/CS)** | | | **log(CIII/CS)** | | | **log(CIV/CS)** | | | **log(CS)** | | |
|---|---|---|---|---|---|---|---|---|---|---|---|---|---|---|---|
| Predictors | *B* | 95% CI | *P*-value (adjusted)[a] | *B* | 95% CI | *P*-value (adjusted)[a] | *B* | 95% CI | *P*-value (adjusted)[a] | *B* | 95% CI | *P*-value (adjusted)[a] | *B* | 95% CI | *P*-value |
| Disease state (PD) | -0.070 | -0.128 – -0.013 | **0.017** (0.068) | -0.043 | -0.102 – 0.016 | 0.148 (0.296) | -0.029 | -0.091 – 0.033 | 0.358 (0.388) | -0.026 | -0.087 – 0.034 | 0.388 (0.388) | 0.028 | -0.098 – 0.153 | 0.663 |
| Age | -0.001 | -0.004 – 0.003 | 0.727 | 0.002 | -0.001– 0.005 | 0.183 | 0.008 | 0.004 – 0.011 | **<0.001** | 0.003 | -2.7e-04 – 0.006 | 0.071 | 3.2e-04 | -0.007 – 0.007 | 0.927 |
| Sex (Male) | 0.018 | -0.036 – 0.071 | 0.510 | 0.021 | -0.034 – 0.075 | 0.455 | 0.008 | -0.048 – 0.064 | 0.785 | 0.026 | -0.030 – 0.082 | 0.351 | -0.047 | -0.164 – 0.069 | 0.420 |
| Smoking | -0.102 | -0.214 – 0.009 | 0.071 | 0.046 | -0.068 – 0.161 | 0.424 | 0.126 | 0.009 – 0.242 | **0.035** | -0.044 | -0.161 – 0.073 | 0.453 | -0.127 | -0.369 – 0.116 | 0.303 |
| Batch 1 | Reference | - | - | - | - | - | - | - | - | - | - | - | - | - | - |
| Batch 2 | 0.047 | -0.048 – 0.142 | 0.325 | 0.163 | 0.066 – 0.260 | **0.001** | 0.088 | -0.003 – 0.179 | 0.057 | 0.057 | -0.043 – 0.156 | 0.259 | -0.037 | -0.243 – 0.170 | 0.724 |
| Batch 3 | -0.125 | -0.216 – -0.034 | **0.008** | 0.113 | 0.019 – 0.207 | **0.019** | -0.242 | -0.329 – -0.155 | **<0.001** | -0.103 | -0.199 – -0.007 | **0.036** | -0.099 | -0.298 – 0.10 | 0.325 |
| Batch 4 | -0.236 | -0.321 – -0.152 | **<0.001** | -0.393 | -0.480 – -0.305 | **<0.001** | -0.307 | -0.386 – -0.229 | **<0.001** | -0.132 | -0.221 – -0.042 | **0.004** | 0.376 | 0.191 – 0.561 | **<0.001** |
| Batch 5 | -0.101 | -0.187 – -0.015 | **0.022** | -0.212 | -0.300 – -0.124 | **<0.001** | - | - | - | -0.204 | -0.294 – -0.113 | **<0.001** | 0.183 | -0.005 – 0.370 | 0.056 |
| Batch 6 | -0.114 | -0.202 – -0.026 | **0.012** | -0.323 | -0.414 – -0.223 | **<0.001** | 0.091 | 0.009 – 0.173 | **0.031** | -0.017 | -0.109 – 0.075 | 0.716 | 0.292 | 0.100 – 0.483 | **0.003** |
| Observations | 87 | | | 87 | | | 71[b] | | | 87 | | | 87 | | |
| $R^2$ / $R^2$ adjusted | 0.447 / 0.382 | | | 0.784 / 0.759 | | | 0.776 / 0.747 | | | 0.379 / 0.306 | | | 0.375 / 0.302 | | |

Abbreviations: CI, complex I activity; CII, complex II activity; CIII, complex III activity; CIV, complex IV activity; CS, citrate synthase activity; x/CS, activity x (CI, CII, CIII, or CIV) normalized to citrate synthase activity; *B*, regression coefficient (unstandardized); 95% CI, 95% confidence interval of the regression coefficient. Significant *P*-values are in bold. Nominal and adjusted *P*-values are given.

[a]Parentheses show *P*-values adjusted for multiple testing using the Benjamini-Hochberg procedure for four tests, i.e., CI/CS, CII/CS, CIII/CS and CIV/CS between the PD and control groups.
[b]Complex III activity measurements from 16 individuals were excluded from the analysis due to technical issues with the reduction of decylubiquinone.

Supplementary Table 6: Linear regression model of enzymatic activity of CI in individuals with PD

| | *Dependent variable* | | |
|---|---|---|---|
| | | **log(CI/CS)** | |
| *Predictors* | *B* | 95% CI | *P*-value |
| Age | 0.001 | -0.005 – 0.006 | 0.763 |
| Sex (Male) | 0.026 | -0.051 – 0.103 | 0.504 |
| UPDRS III score | 0.002 | -0.002 – 0.005 | 0.328 |
| MoCA score[a] | 0.007 | -0.006 – 0.019 | 0.286 |
| Disease duration (months) | -2.0e-04 | -9.0e-04 – 4.0e-04 | 0.518 |
| Batch 1 | Reference | - | - |
| Batch 2 | 0.035 | -0.096 – 0.165 | 0.595 |
| Batch 3 | -0.181 | -0.303 – -0.058 | **0.005** |
| Batch 4 | -0.273 | -0.395 – -0.151 | **<0.001** |
| Batch 5 | -0.119 | -0.236 – -0.001 | **0.048** |
| Batch 6 | -0.128 | -0.235 – -0.020 | **0.021** |
| Observations | 53 | | |
| $R^2$ / $R^2$ adjusted | 0.489 / 0.368 | | |

Abbreviations: CI, complex I activity; CS, citrate synthase activity; CI/CS, complex I activity normalized to citrate synthase activity; *B*, regression coefficient (unstandardized); 95% CI, 95% confidence interval of the regression coefficient; UPDRS III score, sum of MDS-UPDRS part III; MoCA: Montreal Cognitive Assessment; Disease duration (months), duration of motor symptoms in months.
Significant *P*-values are in bold. Nominal *P*-values are given.

[a]Four individuals were omitted from the analysis due to missing MoCA scores.

Supplementary Table 7. Comparison of clinical and demographic characteristics between the PD subgroups

| Variable | CI activity subgroup | Mean ± SD | Median (IQR) | Test | Test Statistic | *df* | *P*-value | Adjusted *P*-value (BH) |
|---|---|---|---|---|---|---|---|---|
| **Age at onset** | Within control range ($n$ = 48) | 61.5 ± 7.8 | 61.1 (12.4) | Student's *t*-test | $t$ = -1.78 | 55 | 0.080 | 0.200 |
| | Below control range ($n$ = 9) | 56.0 ± 11.7 | 54.1 (12.3) | | | | | |
| **MoCA score** | Within control range[a] ($n$ = 44) | 24.8 ± 3.2 | 25 (4) | Student's *t*-test | $t$ = 0.38 | 51 | 0.703 | 0.879 |
| | Below control range ($n$ = 9) | 25.2 ± 3.3 | 25 (3) | | | | | |
| **MDS-UPDRS III score** | Within control range ($n$ = 48) | 29.7 ± 11.5 | 27.0 (15.8) | Wilcoxon rank-sum test | $W$ = 276.5 | NA | 0.189 | 0.315 |
| | Below control range ($n$ = 9) | 23.6 ± 8.8 | 24.0 (9.0) | | | | | |
| **Motor phenotype** | | | | Fisher's exact test | NA | NA | 1 | 1 |
| Within control range ($n$ = 48) | TD/PIGD: 19/21 | | | | | | | |
| Below control range ($n$ = 9) | TD/PIGD: 4/5 | | | | | | | |
| **Sex** | | | | Fisher's exact test | NA | NA | **0.020** | 0.100 |
| Within control range ($n$ = 48) | Male/Female: 33/15 | | | | | | | |
| Below control range ($n$ = 9) | Male/Female: 2/7 | | | | | | | |

Abbreviations: CI, complex I; IQR, interquartile range; *df,* degrees of freedom; BH, Bejamini-Hochberg procedure; Age at onset, Age at onset of motor symptoms; MoCA, Montreal Cognitive Assessment; MDS-UPDRS III, International Parkinson and Movement Disorder Society Unified Parkinson's Disease Rating Scale Part III; TD, tremor dominant; PIGD, postural instability/gait difficulty.

Significant *P*-values are in bold. Nominal and adjusted *P*-values are given. Adjusted *P*-values were calculated using the Benjamini-Hochberg procedure to account for multiple comparisons

[a]MoCA scores were not available from four individuals in the PD group with CI activity within control range.

Supplementary Table 8: Linear mixed effects model of complex I level in single muscle fibers in individuals with PD and controls.

The data are restricted to individuals with available complex I activity data.

| | *Dependent variable* | | |
|---|---|---|---|
| | **log(NDUFB10/VDAC1)** | | |
| Predictors | *B* | 95% CI | *P*-value |
| Disease state (PD) | -0.024 | -0.061 – 0.014 | 0.216 |
| Age | -0.001 | -0.003 – 0.001 | 0.180 |
| Sex (Male) | 0.016 | -0.016 – 0.048 | 0.325 |
| Batch (Batch 2)[a] | -0.096 | -0.130 – -0.062 | **<0.001** |
| **Random Effects** | | | |
| $\sigma^2$ | 0.003 | | |
| $\tau_{00\ \text{Subject}}$ | 0.004 | | |
| ICC | 0.509 | | |
| $n_{\text{Subject}}$ | 62 | | |
| Observations | 6073 | | |
| Marginal $R^2$ / Conditional $R^2$ | 0.253 / 0.634 | | |

Abbreviations: log(NDUFB10/VDAC1), complex I level adjusted for mitochondrial content; *B*, regression coefficient (unstandardized); 95% CI, 95% confidence interval of the regression coefficient; σ2, residual variance; $\tau_{00\ \text{Subject}}$, random intercept variance; ICC, intraclass correlation coefficient, representing the proportion of total variance in the dependent variable attributable to the grouping structure (i.e, subjects); $n_{\text{Subject}}$, number of study subjects.
Significant *P*-values are in bold. Nominal *P*-values are given.

[a]Immunohistochemistry staining was performed in two batches.

Supplementary Table 9. ANOVA comparison evaluating the impact of complex I quantity on complex I activity

| **Model** | **Res.Df** | **RSS** | **Df** | **Sum of Sq** | ***F*** | ***Pr(>F)*** |
|---|---|---|---|---|---|---|
| Model 1 | 57 | 0.64 | NA | NA | NA | NA |
| Model 2 | 58 | 0.76 | -1 | -0.12 | 10.98 | **0.002** |

Comparison of two linear regression models using ANOVA.
Model 1: log(CI/CS)[a]~Disease state + Age + Sex + log(NDUFB10/VDAC1)[b]
Model 2: log(CI/CS)[a]~Disease state + Age + Sex
The table presents the residual degrees of freedom (Res.Df), residual sum of squares (RSS), the difference in degrees of freedom (Df), the difference in residual sum of squares (Sum of Sq), the *F*-statistic (*F*), and the corresponding *P*-value (*Pr(>F)*). Significant *P*-value in bold. Nominal *P*-value is given.
[a]The data have been adjusted for the effect of measurement batch by regressing out this variable.
[b]The data have been adjusted for the effect of staining batch by regressing out this variable.

Supplementary Table 10. Linear regression model of enzymatic activity of CI in individuals with PD and controls.

The data are restricted to individuals with available complex I immunohistochemistry data.

| | *Dependent variable* | | |
|---|---|---|---|
| | | **log(CI/CS)[a]** | |
| *Predictors* | *B* | *95% CI* | *P*-value |
| Disease state (PD) | -0.035 | -0.101 – 0.032 | 0.301 |
| Age | -0.001 | -0.004 – 0.003 | 0.699 |
| Sex (Male) | 0.017 | -0.042 – 0.076 | 0.573 |
| log(NDUFB10/VDAC1)[b] | 0.806 | 0.319 – 1.293 | **0.002** |
| Observations | 62 | | |
| $R^2$ / $R^2$ adjusted | 0.216 / 0.161 | | |

Abbreviations: CI, complex I activity; CS, citrate synthase activity; CI/CS, complex I activity normalized to citrate synthase activity; *B*, regression coefficient (unstandardized); 95% CI, 95% confidence interval of the regression coefficient; log(NDUFB10/VDAC1), complex I level adjusted for mitochondrial content. Significant *P*-values are in bold. Nominal *P*-values are given.

[a]The data have been adjusted for the effect of measurement batch by regressing out this variable.
[b]The data have been adjusted for the effect of staining batch by regressing out this variable.

Supplementary Table 11: Linear mixed effects model of mtDNA copy number in single muscle fibers

| | *Dependent variable* | | |
|---|---|---|---|
| | **mtDNA copy number per micro dissected area (µm$^2$)** | | |
| Predictors | *B* | 95% CI | *P*-value |
| Disease state (PD) | 1.44 | -0.67 – 3.54 | 0.179 |
| Age | 2.2e-03 | -0.11 – 0.11 | 0.968 |
| Sex (Male) | -2.33 | -4.60 – -0.06 | **0.044** |
| Plate 1 | Reference | - | **-** |
| Plate 2 | -5.33 | -7.58 – -3.07 | **<0.001** |
| Plate 3 | -7.22 | -9.38 – -5.07 | **<0.001** |
| Plate 4 | -8.86 | -11.15 – -6.57 | **<0.001** |
| Plate 5 | -5.40 | -8.17 – -2.64 | **<0.001** |
| Plate 6 | -5.95 | -8.70 – -3.20 | **<0.001** |
| Plate 7 | -6.61 | -9.44 – -3.78 | **<0.001** |
| Plate 8 | -6.56 | -9.51 – -3.60 | **<0.001** |
| Plate 9 | -6.55 | -9.22 – -3.88 | **<0.001** |
| Plate 10 | -5.14 | -7.82 – -2.46 | **<0.001** |
| Plate 11 | -4.44 | -7.13 – -1.76 | **0.001** |
| Plate 12 | -5.62 | -8.32 – -2.92 | **<0.001** |
| **Random Effects** | | | |
| $\sigma^2$ | 11.289 | | |
| $\tau_{00\ \text{Subject}}$ | 0.885 | | |
| ICC | 0.073 | | |
| $n_{\text{Subject}}$ | 12 | | |
| Observations | 223 | | |
| Marginal $R^2$ / Conditional $R^2$ | 0.297 / 0.348 | | |

Abbreviations: *B*, regression coefficient (unstandardized); 95% CI, 95% confidence interval of the regression coefficient; Plate, qPCR plate; $\sigma^2$, residual variance; $\tau_{00\ \text{Subject}}$, random intercept variance; ICC, intraclass correlation coefficient, representing the proportion of total variance in the dependent variable attributable to the grouping structure (i.e, subjects); $n_{\text{Subject}}$, number of study subjects.
Significant *P*-values are in bold. Nominal *P*-values are given.

Supplementary Table 12: Linear mixed effects model of mtDNA major arc deletion levels in single muscle fibers

| | *Dependent variable* | | |
|---|---|---|---|
| | **mtDNA deletion fraction** | | |
| Predictors | *B* | 95% CI | *P*-value |
| Disease state (PD) | 0.002 | -0.058 – 0.062 | 0.945 |
| Age | 2.0e-03 | -0.003 – 0.003 | 0.899 |
| Sex (Male) | -0.001 | -0.066 – 0.065 | 0.983 |
| Plate 1 | Reference | - | - |
| Plate 2 | -0.026 | -0.053 – 0.001 | 0.056 |
| Plate 3 | -0.015 | -0.041 – 0.010 | 0.237 |
| Plate 4 | 0.012 | -0.015 – 0.040 | 0.373 |
| Plate 5 | -0.008 | -0.072 – 0.055 | 0.794 |
| Plate 6 | -0.012 | -0.075 – 0.051 | 0.710 |
| Plate 7 | -0.051 | -0.115 – 0.013 | 0.115 |
| Plate 8 | -0.019 | -0.084 – 0.046 | 0.559 |
| Plate 9 | 0.009 | -0.053 – 0.070 | 0.786 |
| Plate 10 | 0.011 | -0.05 1– 0.073 | 0.721 |
| Plate 11 | 5.0e-03 | -0.061 – 0.062 | 0.986 |
| Plate 12 | 0.002 | -0.060 – 0.063 | 0.960 |
| **Random Effects** | | | |
| $\sigma^2$ | 0.002 | | |
| $\tau_{00\ \text{Subject}}$ | 0.001 | | |
| ICC | 0.426 | | |
| $n_{\text{Subject}}$ | 12 | | |
| Observations | 223 | | |
| Marginal $R^2$ / Conditional $R^2$ | 0.094 / 0.479 | | |

Abbreviations: *B*, regression coefficient (unstandardized); 95% CI = 95% confidence interval of the regression coefficient; Plate, qPCR plate; σ2, residual variance; $\tau_{00\ \text{Subject}}$, random intercept variance; ICC, intraclass correlation coefficient, representing the proportion of total variance in the dependent variable attributable to the grouping structure (i.e, subjects); $n_{\text{Subject}}$, number of study subjects.
Significant *P*-values are in bold. Nominal *P*-values are given.

Supplementary Table 13. Linear mixed effects model of heteroplasmic load in single muscle fibers

| | *Dependent variable* | | | | | |
|---|---|---|---|---|---|---|
| | **Heteroplasmic load (amplicon 1)** | | | **Heteroplasmic load (amplicon 2)** | | |
| Predictors | *B* | 95% CI | *P*-value | *B* | 95% CI | *P*-value |
| Disease state (PD) | -3.99 | -15.94 – 7.96 | 0.511 | -6.65 | -24.53 – 11.23 | 0.463 |
| Age | 0.20 | -0.40 – 0.79 | 0.512 | 0.57 | -0.32 – 1.47 | 0.206 |
| Sex (Male) | -3.47 | -15.36 – 8.41 | 0.564 | -8.51 | -26.30 – 9.28 | 0.346 |
| log(NDUFB10/VDAC1) | -22.48 | -67.73 – 22.77 | 0.328 | 40.00 | -18.87 – 98.87 | 0.181 |
| **Random Effects** | | | | | | |
| $\sigma^2$ | 193.48 | | | 289.31 | | |
| $\tau_{00\ \text{Subject}}$ | 43.91 | | | 108.34 | | |
| ICC | 0.18 | | | 0.27 | | |
| $n_{\text{Subject}}$ | 12 | | | 12 | | |
| Observations | 150 | | | 144 | | |
| Marginal $R^2$ / Conditional $R^2$ | 0.060 / 0.234 | | | 0.145 / 0.378 | | |

Abbreviations: heteroplasmic load, the sum of the heteroplasmy levels across the mtDNA region; amplicon 1, rCRS coordinates 600-8,600; amplicon 2, rCRS coordinates 9,100-16,200; log(NDUFB10/VDAC1), complex I level adjusted for mitochondrial content; *B*, regression coefficient (unstandardized); 95% CI = 95% confidence interval of the regression coefficient; σ2, residual variance; $\tau_{00\ \text{Subject}}$, random intercept variance; ICC, intraclass correlation coefficient, representing the proportion of total variance in the dependent variable attributable to the grouping structure (i.e, subjects); $n_{\text{Subject}}$, number of study subjects.
Significant *P*-values are in bold. Nominal *P*-values are given.

Supplementary Table 14. Wilcoxon rank sum tests of mtDNA data in bulk muscle tissue

| **mtDNA copy number per *APP*** | | | | | | |
|---|---|---|---|---|---|---|
| Groups | *n* | Median | IQR | Wilcoxon *W* | *Z* | *P*-value |
| Control | 10 | 10000 | 3215 | 55 | 0.816 | 0.438 |
| Low PD | 9 | 7312 | 3111 | | | |
| Control | 10 | 10000 | 3215 | 38 | -0.178 | 0.894 |
| Normal PD | 8 | 9691 | 16522 | | | |
| Low PD | 9 | 7312 | 3111 | 27 | -0.866 | 0.413 |
| Normal PD | 8 | 9691 | 16522 | | | |

| **mtDNA deletion proportion** | | | | | | |
|---|---|---|---|---|---|---|
| Groups | *N* | Median | IQR | Wilcoxon *W* | *Z* | *P*-value |
| Control | 10 | 0.058 | 0.084 | 34 | -0.898 | 0.391 |
| Low PD | 9 | 0.091 | 0.040 | | | |
| Control | 10 | 0.058 | 0.084 | 43 | 0.267 | 0.824 |
| Normal PD | 8 | 0.072 | 0.066 | | | |
| Low PD | 9 | 0.091 | 0.040 | 46 | 0.962 | 0.361 |
| Normal PD | 8 | 0.072 | 0.066 | | | |

Abbreviations: Normal PD, PD samples with CS normalized CI activity similar to controls; Low PD, PD samples with low CI normalized CI activity.
Significant *P*-values are in bold. Nominal *P*-values are given.

Supplementary Table 15. Linear regression model of heteroplasmic load in bulk muscle tissue as a function of CI activity and enzymatic activity measurement batch.

| | *Dependent variable* | | | | | |
|---|---|---|---|---|---|---|
| | **Heteroplasmic load (amplicon 1)** | | | **Heteroplasmic load (amplicon 2)** | | |
| Predictors | *B* | 95% CI | *P*-value | *B* | 95% CI | *P*-value |
| log(CI/CS) | -2.43 | -14.35 – 9.50 | 0.676 | 8.08 | -7.93 – 24.09 | 0.305 |
| Batch 1 | Reference | - | - | - | - | - |
| Batch 2 | 8.80 | -0.13 – 17.73 | 0.053 | 7.06 | -4.93 – 19.04 | 0.234 |
| Batch 3 | 3.03 | -7.12 – 13.17 | 0.541 | 4.90 | -8.71 – 18.51 | 0.461 |
| Batch 4 | -0.86 | -10.08 – 8.37 | 0.848 | 1.93 | -10.45 – 14.31 | 0.748 |
| Batch 5 | 1.77 | -8.55 – 12.09 | 0.724 | 13.44 | -0.41 – 27.29 | 0.057 |
| Batch 6 | 0.20 | -9.48 – 9.88 | 0.966 | 1.63 | -11.37 – 14.62 | 0.796 |
| Observations | 27 | | | 27 | | |
| $R^2$ / $R^2$ adjusted | 0.220 / -0.014 | | | 0.254 / 0.031 | | |

Abbreviations: heteroplasmic load, the sum of the heteroplasmy levels across the mtDNA region; amplicon 1, rCRS coordinates 600-8,600; amplicon 2, rCRS coordinates 9,100-16,200; CI, complex I activity; CS, citrate synthase activity; CI/CS, complex I activity normalized to citrate synthase activity; Batch, enzymatic activity measurement batch; *B*, regression coefficient (unstandardized); 95% CI, 95% confidence interval of the regression coefficient.
Significant *P*-values are in bold. Nominal *P*-values are given.

Supplementary Table 16. Inclusion and exclusion criteria for the STRAT-PARK cohort

| | |
|---|---|
| Main inclusion criteria for individuals with PD: | 1) Have a diagnosis of clinically established or probable PD according to the movement disorders society (MDS) diagnostic criteria for PD.<br>2) 123I-Ioflupane dopamine transporter imaging (DAT-scan) confirming nigrostriatal degeneration.<br>3) The participant must have the physical and mental capacity to participate in the study.<br>4) The participant must have the capacity to provide written informed consent for study participation.<br>5) Age 20-100. |
| Main exclusion criteria for individuals with PD: | Comorbidity interfering with study participation |
| Main inclusion criteria for controls: | Individuals without neurodegenerative diseases aged 20-100 years |
| Main exclusion criteria for controls: | 1) Diagnosis or clinically established or probable PD, other parkinsonism, or other neurodegenerative disorder<br>2) Clinical suspicion of clinically established or probable PD, other parkinsonism, or other neurodegenerative disorder at screening/baseline<br>3) Comorbidity interfering with study participation |

Supplementary Table 17. Inclusion and exclusion criteria for the NADPARK study

| | |
|---|---|
| Inclusion criteria | 1) A new clinical diagnosis of PD<br>2) Drug naïve state with respect to dopaminergic treatment<br>3) Clinical diagnosis of PD, according to the Movement Disorder Society Clinical Diagnostic Criteria. |
| Exclusion criteria | 1) 123I-Ioflupane dopamine transporter imaging (DAT-scan) without evidence of nigrostriatal degeneration<br>2) Magnetic Resonance Imaging (MRI) suggestive of atypical parkinsonism<br>3) Dementia or other neurological disorder at baseline visit<br>4) Metabolic, neoplastic, or other physically or mentally debilitating disorders at baseline visit. |

Supplementary Table 18. Inclusion and exclusion criteria for the STRAT-COG study

| | |
|---|---|
| Inclusion criteria | 1) Mild dementia according to ICD-11<br>2) A friend or family as a study partner to provide information<br>3) Capacity to provide written informed consent for study participation defined as Montreal Cognitive Assessment (MOCA) score ≥ 16 or Mini Mental State Evaluation (MMSE) score ≥ 20. If there is any doubt regarding the participants capacity to give informed consent we will ask for an independent evaluation by a consultant clinician who is not associated with the STRAT-COG study.<br>4) Age 45-100. |
| Exclusion criteria | Comorbidity that precludes study participation or data interpretation |
| Controls are healthy volunteers and must have no clinical signs of neurodegenerative disease or fulfil the exclusion criteria. Some are partners/spouses of included patients. | |

Supplementary Table 19: Olympus VS120 fluorescence filters

| **Filter** | **Excitation** | **Emission** |
|---|---|---|
| DAPI | 340-380 | 430-480 |
| FITC | 480-500 | 507-543 |
| TRITC | 543-568 | 579-631 |
| CY5 | 630-660 | 669-741 |
